# Stop codon readthrough in *Trichomonas* is a mechanism for gene expression regulation and expanding protein function

**DOI:** 10.64898/2026.08.31.748405

**Authors:** Francisco Callejas-Hernández, Mari Shiratori, Madison Pleas, Steven A. Sullivan, Jane M. Carlton

## Abstract

*Trichomonas vaginalis* is the causative agent of trichomoniasis, a common sexually transmitted infection among women of reproductive and peri-menopausal age. The parasite has an unusually large genome, rich in complex repeats, including a vast repertoire of transposable elements and multi-copy gene families. Since very few *T. vaginalis* genes have introns, gene expression is usually straightforward, with ribosomal translational machinery proceeding from a start codon to the next in-frame stop codon of an unspliced poly(A)denylated mRNA. However, our previous studies raised the possibility of *T. vaginalis* gene expression involving stop codon readthrough (SCR), where transcription through in-frame stop codons produces longer-than-predicted mRNAs that translate to fully functional proteins. Here, we leverage long-read RNA-seq and new chromosome-scale assemblies of two *T. vaginalis* strains and two avian sister species to investigate and characterize ∼1,400 long, mature mRNAs that contain more than one predicted protein-coding gene transcribed from what we call ‘RT genes’, composites of adjacent predicted genes. We first identify RT genes in a second *T. vaginalis* strain and in close relatives *T. vaginalis-like* and *T. stableri*, indicating that this phenomenon is conserved among *Trichomonas* species and strains. Second, we find transcripts of RT genes to be more abundant by many orders of magnitude than monocistronic genes. Third, we found the distance between predicted genes within RT genes to be significantly shorter than between adjacent independent predicted genes. Fourth, functional annotation revealed that RT genes encode at least 50 distinct protein functions, suggesting that this unusual transcriptional mechanism has a role in an array of biological processes in *Trichomonas*. Our results from two *Trichomonas* species suggest that SCR is an important mechanism controlling gene expression and the diversity of protein function in this parasite.

## Introduction

Species of *Trichomonas,* phylum Parabasalia, are unicellular, eukaryotic, flagellated, extracellular parasites that inhabit the digestive, respiratory, and urogenital tracts of animals and humans. One species, *Trichomonas vaginalis,* is a human urogenital parasite that causes trichomoniasis, a common sexually transmitted infection affecting millions each year (1, 2). The female reproductive tract is an especially dynamic niche, with fluctuating iron levels, pH, microbial communities, oxygen stress, and immune responses that the parasite must contend with to establish an infection (3). The parasite must rapidly modulate its gene expression to survive in such diverse and changing host environments.

*T. vaginalis* exhibits a simplified and unusual gene expression system compared with other protozoans. Unlike intron-rich eukaryotic parasites such as *Plasmodium* or *Toxoplasma*, *Trichomonas* has only a small number of genes with spliceosomal introns, suggesting extensive intron loss during evolution (4). Gene regulation appears to rely heavily on transcriptional flexibility, RNA processing, and post-transcriptional mechanisms rather than extensive promoter complexity or alternative splicing (5–7).

Recent advances have further increased our understanding of *T. vaginalis* gene expression. Muñoz *et al.* (2025) reported that upon direct contact with host epithelial cells, chromatin accessibility increases at specific promoters, correlating with upregulation of key pathogenesis genes, including BspA-like adhesins, Bap-like proteins, cysteine proteases, and serine/threonine kinases. This chromatin remodeling enables the parasite to switch from a free-swimming metabolic program to one optimized for adhesion and cytotoxicity (8). However, more evidence is needed to show that changes in promoter accessibility drive the transcriptional response. Parasite-to-parasite communication via *T. vaginalis* extracellular vesicles (TvEVs) also plays a role in regulating parasite gene expression. TvEVs are released and communicate with neighboring parasites, transferring virulence factors that increase the parasite’s adherence capacity up to ∼3-fold and enhance its ability to colonize host cells (9). The vaginal microbiome has also been shown to influence *T. vaginalis* gene expression; for example, exposure to the vaginal early colonizer *Bifidobacterium breve* temporarily boosts *T. vaginalis* growth and reshapes its lipid metabolism through direct microbiome-to-parasite gene regulation (10). Finally, certain *T. vaginalis* strains carry an endosymbiotic dsRNA Trichomonas virus (TVV), and cells with higher viral loads exhibit distinct levels of stress, adhesion, and chromatin-related gene expression (11).

During the project to sequence the first *T. vaginalis* genome in 2007 (12), we observed, based upon high sequence similarity to known proteins in public databases, that a total of 1,465 *T. vaginalis* genes appeared to be ‘split’, i.e., the N-terminal and C-terminal ends of the protein were encoded by adjacent predicted genes. Of these split genes, 923 were caused by a frameshift, and 542 were caused by a point mutation resulting in a premature stop codon. It was unclear whether these represented recent pseudogenes, since many appeared to have intact matches elsewhere in the genome, or whether they suggested a novel translational frameshifting mechanism in *T. vaginalis* (12). Subsequently, in 2012, Kay *et al*. analyzed *T. vaginalis* ATP-binding cassette (ABC) membrane transport genes in the published genome and showed that *T. vaginalis* likely can read through in-frame stop codons of ‘partial’ ABC genes to generate fully functional proteins (13). The authors concluded that this suppression of stop codons by *T. vaginalis* warranted further investigation of its mechanism and frequency. Here, using our newly assembled chromosome-scale genomes and high-fidelity long-read RNA-seq datasets from four *Trichomonas* species and strains infecting humans and birds, we confirm and further characterize such “readthrough (RT) genes”. We found that this unusual mechanism of gene expression is conserved among *Trichomonas* species we analyzed, and that RT genes code for proteins with a wide range of functions. We speculate that the ability to read through stop codons boosts the parasite’s ability to add new sections of genes that code for protein domains or regulatory signals and allows for expanded protein functions without the need to generate new genes.

## Material and methods

### Parasite culture and long-read RNA-seq library preparation

*T. vaginalis* laboratory strains G3 (14) and MOR31 (15) were cultured in Diamond’s Tryptose-Yeast-Mannose (TYM) media adjusted to pH 6.2. *Trichomonas vaginalis*-like strain 3688 isolated from a mourning dove in Arizona, and *Trichomonas stableri* strain BTPI-3 isolated from a Pacific coast band-tailed pigeon in California (16), were cultured in Hollander’s Fluid media adjusted to pH 6.8. Both media were supplemented with 10% horse serum (ThermoFisher Scientific, #26050088), 1% penicillin and streptomycin (Invitrogen, #15140-122), and an iron solution comprised of ferrous ammonium sulfate and sulfosalicylic acid. *T. vaginalis*-like 3688 and *T. stableri* BTPI-3 were treated with the Mycoplasma Removal Agent (MRA, BioRad, #BUF035) for 1 week and were confirmed negative by a *Mycoplasma* detection PCR protocol. *T. vaginalis* strains G3 and MOR31 were treated previously and determined to be *Mycoplasma*-free.

Log-growth-phase cultures of each of the parasite species were prepared in triplicate, and 5 million parasites per sample were lysed in RLT lysis buffer containing 1% 2-mercaptoethanol. Total RNA was isolated using the Qiagen RNeasy Mini Kit (#74104), including a column DNase treatment using the Qiagen RNase-free DNase kit (#79254). RNA concentration and quality were quantified using a Nanodrop One spectrophotometer and Agilent TapeStation 4200. All RNA samples had an RNA Integrity Number (RIN) value > 8. RNA samples were sent to Maryland Genomics for PacBio Kinnex long-read RNA library preparation. Libraries were sequenced using a PacBio Revio SMRT Cell Sequencing run (HiFi/CCS mode) for 24 hours.

### Long-read RNA-seq processing and transcript assembly

Sequencing reads were mapped to their respective reference genome sequences: *T. vaginalis* strain G3 (2022 new assembly and annotation; JAOSJJ000000000.2) (17); *T. vaginalis* strain MOR31; *T. vaginalis*-like strain 3688; and *T. stableri* strain BTPI-3 (17), using minimap2 (v 2.3) in splicing-aware mode, with secondary alignments not allowed. Remaining chimeric alignments in the BAM files were removed as described in Callejas-Hernandez *et al.* (4).

RT gene candidates were identified following a series of filtering steps and manual curation (**Figure 1**). After transcript assembly using StringTie (v2.2.1), transcripts that overlapped entirely or partially with more than two predicted genes in the three RNA-seq replicates for each strain were considered the first candidates. A second filter was applied based on the coverage of long reads supporting the transcript length. Only transcripts whose length was supported by long reads of at least 80% of the total transcript length survived the filtering. Next, poly(A) tails at the ends of transcripts were identified using a custom Python script. Poly(A) tails (minimum eight nucleotides, with up to one mismatch) at the end of soft-masked alignments with a minimum coverage of 10 reads were considered *bona fide* poly(A) signals. Candidate transcripts were classified based on the number of long reads supporting at least 80% of the total transcript length and the presence of poly(A) tails as follows: (1) no-confidence (zero long reads support the transcript length); (2) low-confidence (support from one or two replicates); and (3) high-confidence (support from three replicates with a minimum coverage of 10, and presence of poly(A) tails at the transcript end). Transcript coordinates from the three RNA-seq replicates were merged, extending the coordinates to the longest transcript. Finally, all candidates were manually curated based on the three parameters listed above and the coverage profile.

**Figure 1.**
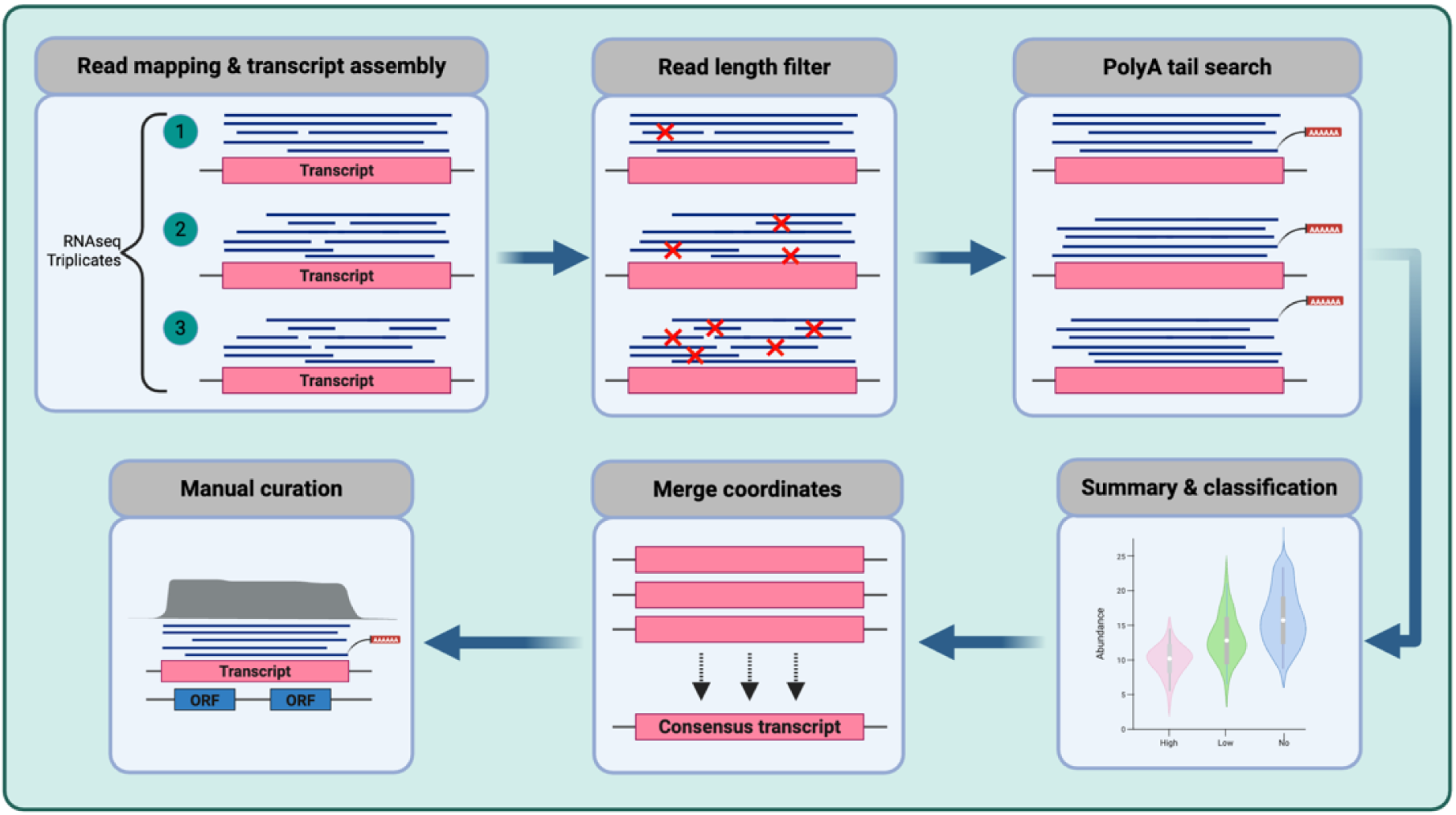
Transcriptome analysis and RT gene identification. Transcripts were mapped and filtered according to length (80% coverage) and 3’ poly(A) tails, assembled, and manually curated to identify RT genes.

### Substantiation and functional annotation of RT gene candidates

The intergenic distance (sequence length separating two adjacent genes on a chromosome) and transcript abundance (Transcripts per Million; TPM) of the RT genes were calculated and compared to the genome-wide transcriptome using a non-parametric Mann-Whitney U test. To reduce the imbalance effect (∼100 vs 17,000 samples), the analysis was repeated 1,000 times using random sampling (balanced sample sizes) to summarize the effect size and p-value distributions (Hodges-Lehmann estimates). Only genes with a TPM higher than 1 were considered for analysis.

Orthologs and paralogs were identified using a combination of MCL and BLASTN, with custom Python scripts and minimum cutoff thresholds of e-value 0.001, percent identity 90%, and minimum query coverage of 50%.

Functional annotation of the RT gene candidates was investigated using the current genome annotation of *T. vaginalis* G3. The putative functions were classified as eukaryotic-associated, bacterial-associated, or both (universal) based on the obtained product name (predicted functions).

## Results

### Long-read RNA-seq reveals transcripts encompassing >1 predicted gene

Long-read RNA-seq libraries were generated in triplicate for four different strains of *Trichomonas:* strains G3 and MOR31 of *T. vaginalis* isolated from humans, and two different species found in birds, *T. vaginalis-*like 3688 and *T. stableri* BTPI-3, the closest known relatives to *T. vaginalis* (17). An average of 6.7 million HiFi PacBio long-read RNA-seq reads per triplicate sample was obtained, with an average read length of 1.7 kb, in accord with the expected average gene length (1.6 kb), and a mean quality of 25.70 (**Supplementary Table 1**). The mapping rate to all reference genomes exceeded 90% for all samples.

Given the availability of a highly curated genome and annotation of *T. vaginalis* G3, we analyzed its transcriptome first. We identified 1,497 long transcripts encompassing more than one predicted protein-coding gene, and named these “RT gene candidates” (**Figure 2A, B**), 496 of which were further classified as ‘high confidence’ by read length and PAS filters (**Figure 1**). Approximately 90% of these high-confidence candidate transcripts overlap only two predicted protein-coding genes and are consistently shorter than those classified as low- or no-confidence. Transcripts lacking poly(A) tails, or whose length was not supported by long reads, are longer and overlap up to eight predicted genes, suggesting that they are low-quality sequences or artifacts of faulty mapping or assembly. The complete set of candidates and their coordinates is described in **Supplementary Table 2**.

**Figure 2.**
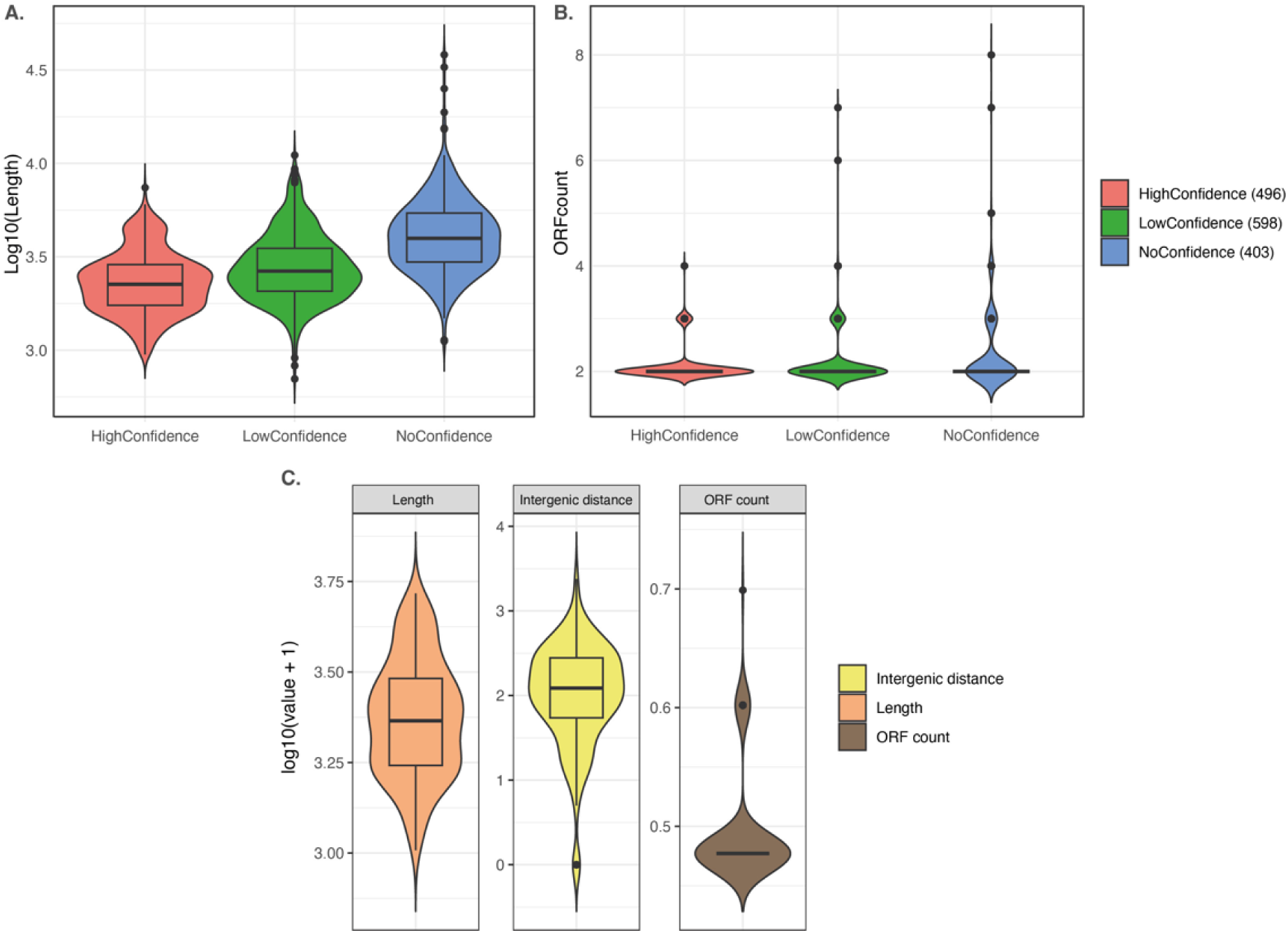
*T. vaginalis* G3 RT gene candidate characterization. (A) Gene length of RT gene candidates classified as high, low, and no confidence. (B) Number of predicted genes within the RT gene candidates. (C) Intergenic distance, length, and gene count of the final curated 118 RT gene candidates.

After extensive manual curation, 118 *T. vaginalis* G3 RT genes (**Figure 2C**) were used as the final set for further analyses. These RT genes have a mean length of 2.5 kb, ∼1 kb longer than the transcriptome mean read length, and the predicted genes they encompass have a mean intergenic distance of 204 bp (range: 0 - 2.3 kb). Almost all candidates encompass just two predicted genes; 16 overlap three, and one overlaps four (**Figure 2C**).

**Figure 3A** shows an example of an RT gene (candidate-1371) supported by strong long-read RNA-seq evidence, uniform coverage, and 3’ poly(A) signals across triplicate RNA-seq samples. The first predicted gene in candidate-1371 (TVAGG3_0345120) encodes a protein involved in signal transduction (phosphoinositide-interacting signal transduction regulators), and the second (TVAGG3_0345110) encodes a protein with copies of the ankyrin repeat, which is thought to be involved in protein-protein interactions and is the most abundant repeat motif in eukaryotic proteins. **Figure 3B** is a second example of an RT gene (candidate-823) that encompasses three predicted protein-coding genes (TVAGG3_0059270, TVAGG3_0059280, TVAGG3_0059290) encoding the halves of an ATP-binding cassette (ABC, group A) protein previously described by Kay *et al*., 2012 (13). Consistent with their findings, functional annotation of the encompassed genes describes the motifs needed to produce a fully functional ABC protein. Within our 118 high-confidence manually curated RT genes, we identified two of the three previously described SCR events reported by Kay *et al* for ABC genes as RT gene candidate-823 and candidate-361. Although the two adjacent genes (TVAGG3_0372580, TVAGG3_0372590) reported in the third previously described SCR event are conserved with 100% identity in the new chromosome-level assembly, no long-read RNAseq supporting a transcript for either of the two predicted genes was found mapped to that locus.

**Figure 3.**
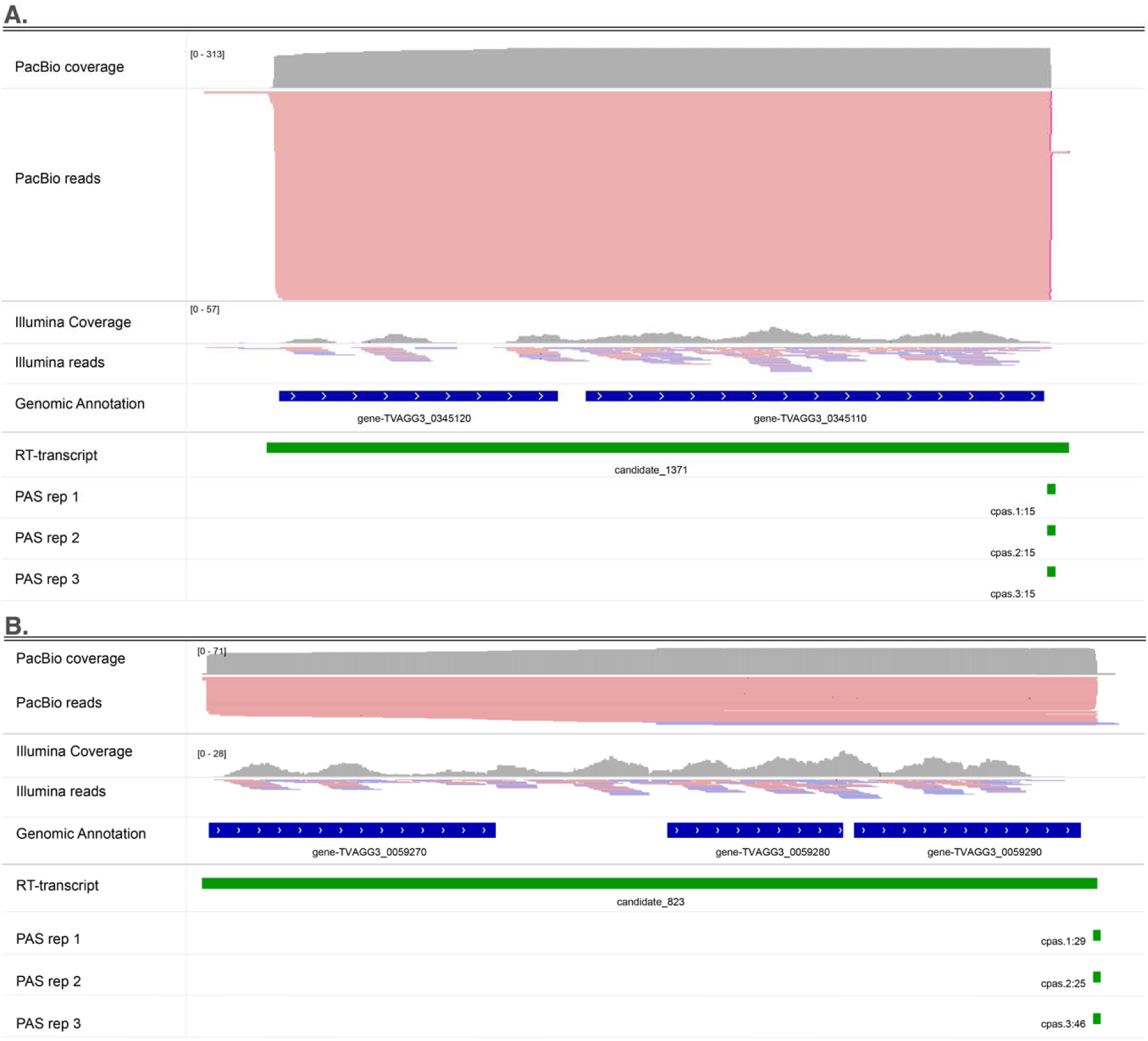
RT gene IGV screenshots of two examples of RT genes: (**A**) Candidate-1371, a newly identified RT gene with high, uniform long-read coverage (PacBio) and poly(A) signals (PAS) at the 3’ end, supported in triplicate. (**B)** Candidate-823, a transcript identified by Kay et al. (13) and confirmed here that overlaps three genes encoding an ABC transporter gene.

The length, standard intergenic distance (between adjacent genes within RT genes), and transcript abundance of the 118 manually curated RT genes were compared to the same statistics calculated for the genome-wide transcriptome (118 RT genes vs ∼17,000 genes; **Figure 4**. First, our results show that RT genes are, on average, significantly longer than the rest of the transcripts in the transcriptome (**Figure 4A**). The mean length of RT genes is 2.5 kb (4.06 kb at the 90th percentile) versus 1.6 kb for all the transcripts (2.9 kb at the 90th percentile). Balanced random subsampling analysis also confirmed that the average length is significantly different (pvalue < 0.001) between the two groups (**Figure 4B**). Second, the mean intergenic distance of the genes within RT genes is 186 bp (median 106 bp) with a 90^th^ percentile of 405 bp versus a 7 kb mean (median 900 bp) and 90^th^ percentile of 22.97 kb for the rest of the single gene-containing transcripts (**Figure 4C**). In other words, genes within RT genes are closer to each other than other non-RT genes across the genome. Results of 1000 random comparisons also confirmed this finding (**Figure 4D**).

**Figure 4.**
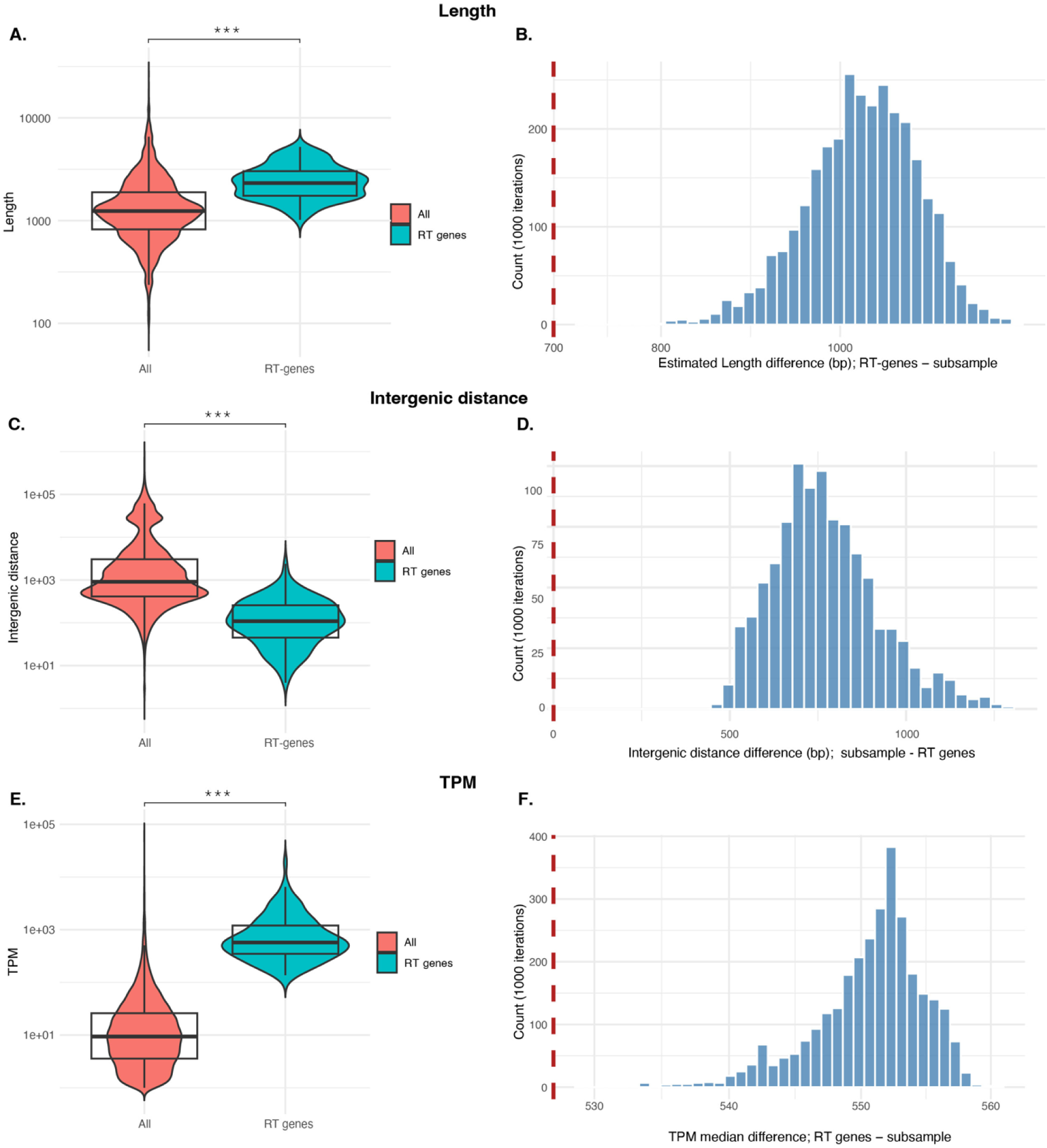
Statistical analysis of the transcript length, intergenic distance, and transcript abundance of RT genes compared to the whole transcriptome. **A, C, and E** show transcript length, intergenic distance, and TPM distributions, respectively, of the RT genes compared to the whole transcriptome. **B, D, and F** show the distribution of Hodges-Lehmann estimates over 1000 random comparisons for transcript length, intergenic distance, and TPM distributions, respectively. The vertical dashed red line indicates the point of no difference (zero median shift) between the two groups.

The third comparison of transcript abundance of RT genes versus the rest of the transcriptome showed RT gene transcripts are significantly more abundant (**Figure 4E, F**). The mean TPM for RT genes is 1,136 (2,757 at the 90^th^ percentile), whereas for the remaining transcripts, the mean TPM is 50.1 (67.7 at the 90^th^ percentile; **Figure 4E**). Results of 1000 random comparisons also confirmed this finding (**Figure 4F**) with a pvalue < 0.001.

The predicted functions of genes within the 118 RT transcripts were analyzed by scanning against the Interpro protein signature database, returning Interpro matches for 94 of them (**Figure 5**). In 46 of the RT transcripts, the genes share the same or related predicted function, whereas in 48 transcripts, the genes have a mix of predicted functions that are biologically unrelated (the remaining genes were considered hypothetical proteins of unknown function). In both cases, the most abundant predicted protein functions correspond to the most abundant predicted functions in the *T. vaginalis* proteome, such as BspA, ABC transporters, kinases, Armadillo repeats, and ankyrin repeats (**Figure 5A**). These five functions are represented as the most expanded multi-copy gene families in *T. vaginalis*, although only a small proportion of multi-copy families were represented among the RT genes (**Figure 5C**). Combined, a total of 53 protein functions were coded for by the 118 RT genes, and their classification as eukaryotic-associated, bacterial-associated, or both (universal) revealed that 17% (9 genes) have a potential bacterial origin (**Figure 5B and Supplementary Table 3**).

**Figure 5.**
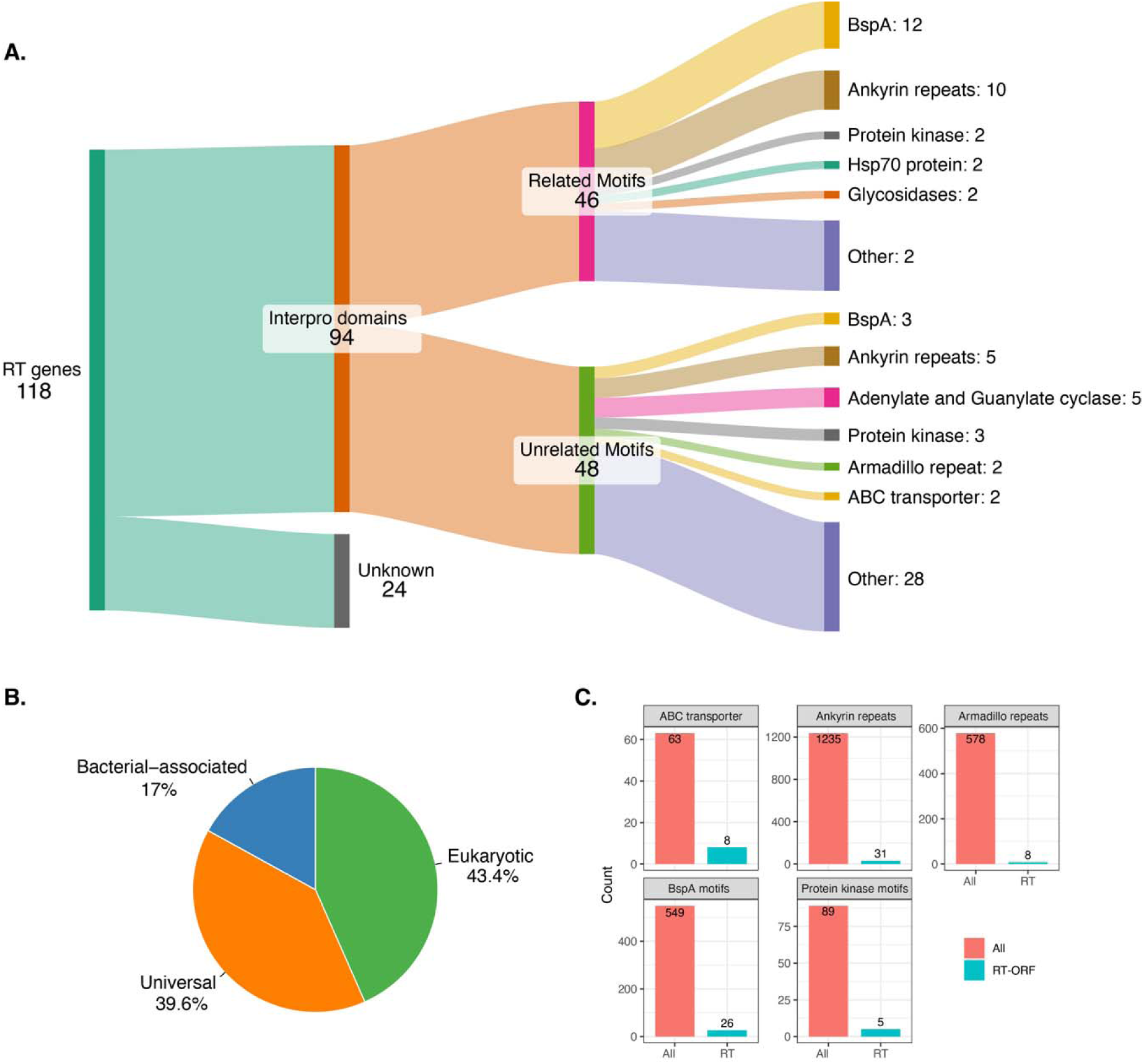
Functional annotation of proteins encoded by RT genes. **(A)** A Sankey plot summarizing RT genes with Interpro hits (related motifs = genes within the same RT gene with the same or related motifs; unrelated motifs = genes within the same RT gene with unrelated/mixed motifs). **(B)** RT genes with predicted functions (94) classified as strictly eukaryotic, bacterial-associated, or both (universal). (**C)** The top abundant gene families in *T. vaginalis* and the total present within the RT genes.

### Stop codon readthrough evidence is found in closely related *Trichomonas* strains and species

We expanded the search for RT genes using long-read RNAseq to a second *T. vaginalis* strain, MOR31, and two species isolated from birds, *Trichomonas stableri* strain BTPI-3, and *Trichomonas vaginalis*-like strain 3688 (**Table 1**), that are closely related to *T. vaginalis*. Only RT gene candidates classified as high confidence (for the variables described in **Figure 1**) were selected for further analysis. *T. vaginalis* MOR31 had a larger RT gene repertoire than strain G3, with 448 candidates, while *T. vaginalis*-like 3688 had 188, and *T. stableri* BTPI3 had 126. For some of those candidates, multiple copies (paralogs) were identified within a 90% nucleotide identity threshold.

**Table 1.** Summary of RT genes found in several *Trichomonas* species and strains. Some RT genes were identified as multi-copy genes (90% nucleotide similarity threshold).

| Species | RT genes | Multi-copy | Singletons |
| --- | --- | --- | --- |
| *T. vaginalis* G3 | 118 | 0 | 118 |
| *T. vaginalis* MOR31 | 448 | 43 | 405 |
| *T. stableri* BTPI-3 | 126 | 12 | 114 |
| *T. vaginalis*-like3688 | 188 | 7 | 181 |

We compared the statistics (number of predicted genes per RT gene, RT gene length, intergenic distance between adjacent genes within RT genes, and transcript abundance relative to the whole transcriptome) of the RT genes in these additional *Trichomonas* species and strains to *T. vaginalis* G3 (**Figure 6**). *T. vaginalis* MOR31 and *T. vaginalis*-like 3688 have RT genes containing up to four predicted protein-coding genes, with two predicted genes per RT gene being most common (a similar distribution in *T. vaginalis* G3). In contrast, all the RT genes in *T. stableri* BTPI-3 contain only two predicted protein-coding genes (**Figure 6A**). In agreement with the results for *T. vaginalis* G3, RT genes in the other trichomonads have a mean length of 2.3 kb and are significantly longer than the rest of the transcriptome (mean length ∼1.5 kb) (**Figure 6B**). Although the intergenic distance among non-RT genes is shorter in *T. stableri* BTPI-3 and *T.vaginalis*-like 3688 (mean ∼1.6 kb) than in *T. vaginalis* strains G3 and MOR31 (∼6 Kb), it is still significantly longer than among RT genes, which is consistently short (∼250 bp) among all the strains and species analyzed (**Figure 6C**). Our results also confirmed that RT gene transcripts are more abundant (mean TPM >1,000) than the rest of the transcripts (mean TPM ∼50) by several orders of magnitude (**Figure 6D**). Lastly, a nucleotide-level conservation analysis revealed that a small proportion of RT genes are conserved across strains and species. In agreement with our hypothesis that stop codon readthrough is a relevant gene expression mechanism in *Trichomonas*, we found high conservation levels of some of the candidates across species. *T. vaginalis* strains MOR31 and G3 were found to share the largest number of RT genes, with a total of 59. With the largest repertoire (448), MOR31 also shares RT genes with *T. vaginalis*-like (22). The remaining genome pairs share only one to six RT genes (**Figure 6E**).

**Figure 6.**
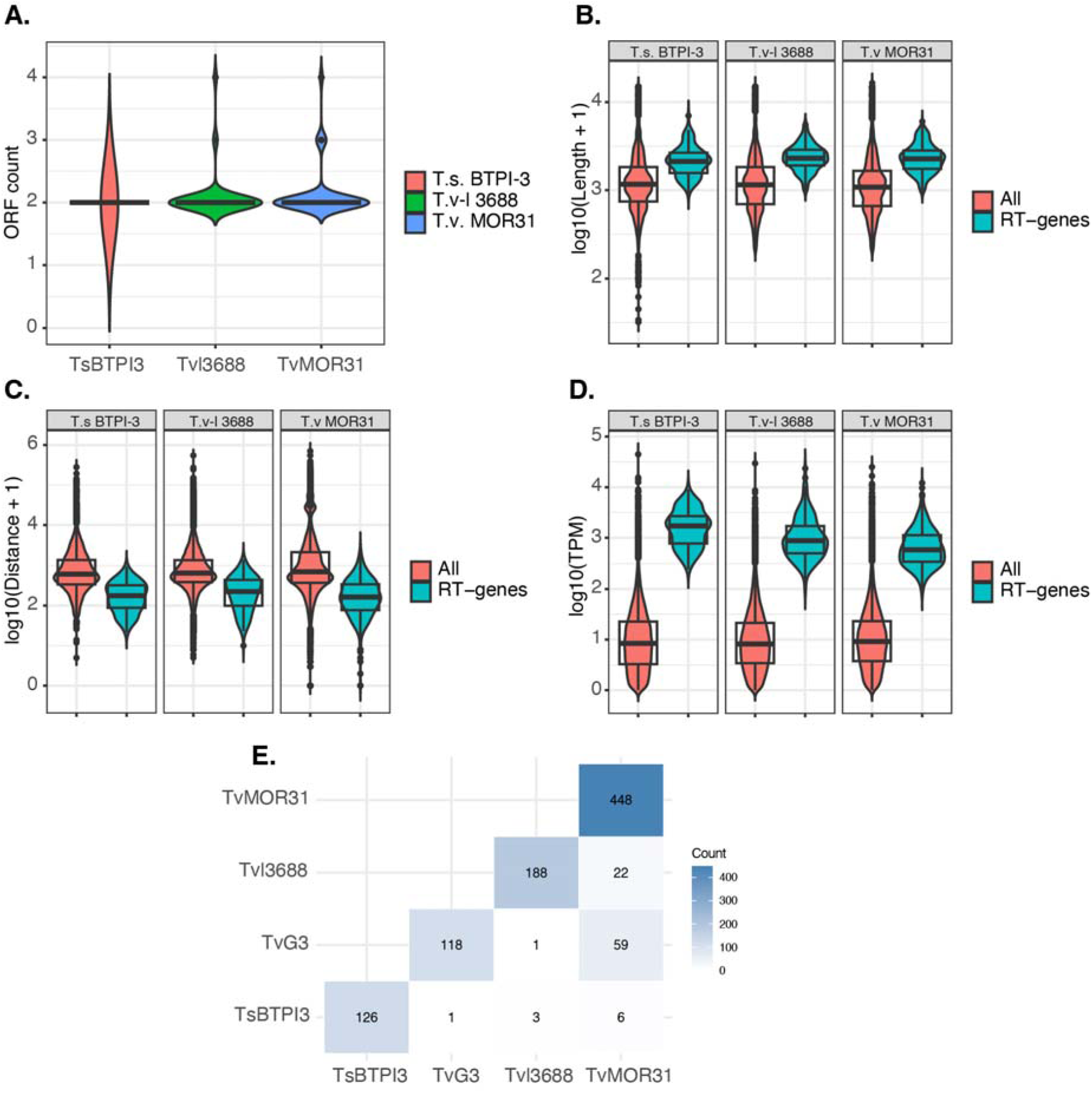
Summary of the RT genes identified in three other *Trichomonas* species and strains. **(A)** Predicted protein-coding gene count per RT gene. **(B)** RT-gene length distribution. (**C**) RT gene intergenic distance**. (D**) RT gene transcript abundance**. (E)** Heat map showing the number of RT genes conserved between species and strains (only the lower matrix is shown). The y-axis of B, C, and D uses a log10 scale for visualization purposes.

Among the 59 conserved RT genes between G3 and MOR31, a total of 30 different putative protein functions were identified, including those from the most expanded multi-copy gene families and similar frequency as described in **Figure 5A**.

The one RT gene found in all four parasite genomes is ∼2 kb long, and its constituent predicted genes encode two proteins of unknown function. This RT gene is highly conserved, with percent similarity ranging from ∼99% (*T. vaginalis* strains G3 and MOR31) to ∼97% between *T. vaginalis* strains and *T. vaginalis*-like. The RT gene from the bird-infecting species (*T. stableri* BTPI-3) is the least conserved, with 94% similarity, and it also has the longest 5’ UTR (**Figure 7**). Although the four sequences contain several SNPs, both predicted genes are conserved, and, interestingly, the ‘intergenic’ region is the most conserved, with only one SNP in the *T. stableri* sequence.

**Figure 7.**
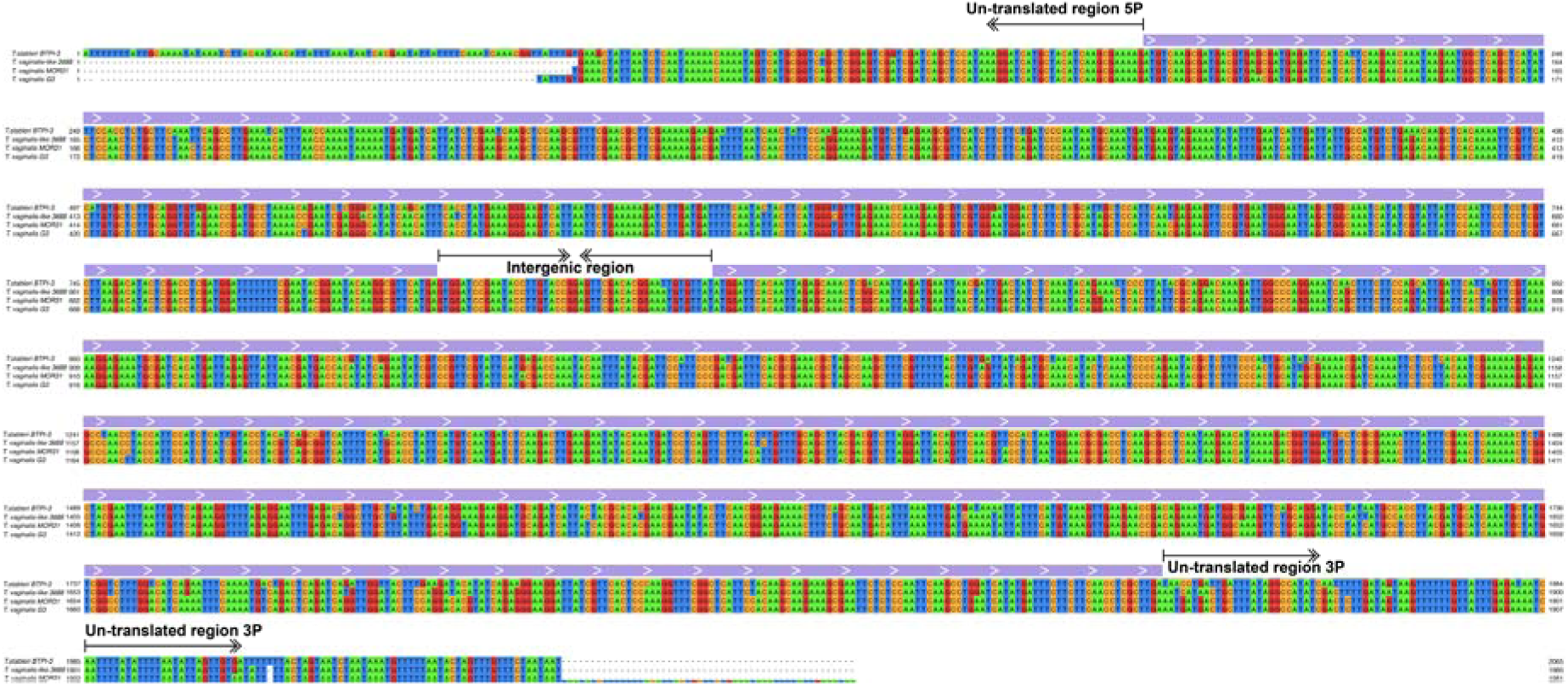
Multiple sequence alignment of the single RT gene conserved among the four parasite genomes analyzed. Nucleotides are represented in different colors, and the Predicted genes are delimited with the upper violet block above the alignment. The 5’ UTRs, 3’ UTRs, and intergenic regions are indicated by black arrows.

## Discussion

Stop codon readthrough (SCR) is a biological phenomenon whereby ribosomes ‘read through’ a stop codon and continue translating until it reaches another stop codon (typically the next in-frame stop codon).

We divide eukaryotic SCR into two main categories by analyzing its overall characteristics across different species: The first category includes SCR in higher eukaryotes that is promoted by stress conditions, resulting in aberrant transcripts that are not necessarily translated. For instance, in humans, stress conditions such as viral infection can derange transcription termination, leading to aberrantly longer (or shorter) proteins. Such aberrant SCR transcripts are sometimes referred to as “downstream of a gene” (DoGs) and are, in most cases, depleted of strong poly(A)denylation signals (PAS) and retained in the nucleus, and thus not translated (18, 19). To our knowledge, a similar mechanism has been described in only two protozoan species. Shalev-Benami *et al.,* (20) demonstrated that the aminoglycoside antibiotic paromomycin induces dose-dependent UGA stop codon readthrough (reaching ∼26% misreading at IC₅₀) in an *in vitro Leishmania* translation system, resulting in C-terminally extended proteins and correlating with inhibition of parasite growth. This suggested that SCR events in trypanosomatids do not occur naturally and that the loss of ribosomal decoding fidelity is detrimental to the parasite. Second, a comprehensive genome-wide translation analysis of *P. falciparum* asexual blood stages using ribosome profiling provided evidence that stop codon readthrough occurs in a few (∼15) genes, and only extends a short way into the nominal 3’ UTR (21). The biological effect of these extended proteins is unknown.

The second category of SCR involves reading through more than one predicted gene. It is proposed to represent a major regulatory mechanism for generating proteome diversity in species of Diptera (flies and mosquitoes). Jungreis *et al.,* (22) identified ∼600 SCR events (what we call RT genes) in 21 species of *Anopheles* mosquitoes and >900 in 20 *Drosophila* species, an average of ∼5% of the total transcriptome in both species. In all cases, RT genes spanned two or more predicted genes and were naturally expressed (23). Their analyses also identified downstream predicted genes exhibiting strong evolutionary signatures of protein-coding selection (e.g., conserved amino acid changes, low Ka/Ks ratios) and provided evidence for readthrough as a functional, often regulatory, mechanism of gene expression. For instance, they found that SCR determines peroxisomal targeting of the final protein, and that the extension adds functional domains or regulatory signals to the protein. Intriguingly, ribosome profiling in *Drosophila* showed higher translation efficiency of RT gene transcripts than monocistronic transcripts (23). Why SCR exists as an alternative gene expression mechanism in flies, when cis- and trans-splicing mechanisms also operate in them, is not known. However, SCR is hypothesized to enable the rapid addition of new protein domains or regulatory signals without requiring new genes, acting as an “evolutionary catalyst” (22).

Our SCR results in Trichomonads fall into the second category. We first described “split” genes in *T. vaginalis* during annotation of the reference G3 genome sequence (12). Subsequently, during analysis of the repertoire of ATP-Binding Cassette (ABC) transporter genes in the *T. vaginalis* draft genome, Kay *et al.* identified three SCR events (13). They found that none of the 98 *T. vaginalis* ABC genes alone contained all the motifs required to encode a full-length protein, instead encoding “half-transporters”, where the halves encoded by adjacent predicted genes formed a complete protein. They presented RT-PCR and EST evidence for three of these genes, indicating that *T. vaginalis* can read through TGA stop codons to translate full-length ABC proteins. Here, we identified 1,497 long transcripts that completely encompass more than one predicted gene. A third of these were validated as present in three biological replicate datasets and as full poly (A)-tailed transcripts. After a series of additional stringent filtering steps to reduce potential false positives, we defined a final repertoire of 118 “high-confidence” transcripts as read-through (RT) genes. Among this set were two of the three RT gene candidates (361 and 823 in our numbering) described by Kay *et al.* (13). Their third RT gene candidate (encompassing TVAG_072410 and TVAG_072420) lacked long-read RNA-seq evidence supporting an RT gene in our data.

Our 118 *T. vaginalis* G3 RT genes display several key characteristics. First, these transcripts are much longer and far more abundant than monocistronic transcripts. Second, the distances between predicted genes within RT genes were much shorter than the intergenic distances between monocistronic genes, suggesting that close gene proximity is essential for SCR. Third, there was a nearly even distribution of TAA, TGA, and TAG stop codons across the 118 RT genes, suggesting an absence of ‘leaky’ stop codons. Fourth, the RT genes encode proteins involved in a range of functions, including ABC transporters, and comprise members of 53 different gene families, implicating RT genes in almost all biological processes in *T. vaginalis*.

These observations suggested that SCR readthrough might be an important mechanism of gene expression among Trichomonads. Our analyses of additional *Trichomonas* species and strains, comprising *T. vaginalis* strain MOR31 and two bird-infecting species, *T. vaginalis*-like 3688 and *T. stableri* BTPI-3, support that idea. We confirmed the presence of RT genes in these species, and many of their characteristics compared with monocistronic transcripts were conserved across genomes, including longer transcript lengths, higher abundance, and shorter intergenic distances. In addition, the number of predicted genes encompassed by RT genes (two to four) is similar among the trichomonads we examined, with the predicted genes encoding a broad range of putative protein functions. Notably, we found some RT genes to be highly conserved between strains.

Polycistronic transcription is the only known mechanism in protozoan parasites for generating long RNAs containing multiple predicted genes. It has long been known that trypanosomatids such as *Trypanosoma brucei*, *Trypanosoma cruzi, and Leishmania,* which cause African sleeping sickness, Chagas disease, and Leishmaniasis, respectively, use this mechanism as the first step for gene expression (24). Long pre-mRNAs referred to as polycistronic transcription units (PTUs) containing several not necessarily functionally related predicted genes, are trans-spliced using a conserved mini-exon (aka ‘splice leader’ (SL)), to yield monocistronic transcripts ready to mature at both ends (5’ 7-methylguanosine cap and a 3’ poly(A) tail) and be translated as independent proteins (24). Although PTUs are not translated (as such), some of them and their trans-splicing sites have been identified in trypanosomatids along with their conserved motifs required for their processing by trans-splicing, which differentiate them from bacterial operon-like structures (25–27).

A recent study of *Cryptosporidium parvum* identified polycistronic transcripts with up to four predicted genes coexisting with transcripts of their corresponding monocistronic genes in at least ∼10% of the parasite’s genes (28). In the absence of genes encoding trans-splicing machinery and trans-splicing leader sequences, the authors hypothesized that these genes are controlled by a trans-splicing-independent system, using either alternative start sites or RNA processing mechanisms. Trans-splicing has also been identified in the luminal parasite *Giardia lamblia* (another anaerobic parasitic protist) (29), although it is limited to only four genes and does not involve polycistrons. While a recent study reported that *T. vaginalis* may be capable of exon processing by trans-splicing (30), we did not find evidence that RT genes can be translated or processed by this or a similar mechanism. In our final set of 118 *T. vaginalis* G3 RT gene candidates, no poly(A) signals were found within the transcript coordinates (including within the intergenic region), casting doubt on the possibility that a polycistronic mechanism translates these transcripts into independent proteins.

To conclude, our data suggest that ribosomes in several species/strains of *Trichomonas* can read through stop codons, a biological phenomenon described in viruses, ciliates, mosquitoes, and higher eukaryotes (31–35). The mechanism of trichomonad SCR is yet to be determined, but possibilities include leaky stop codons, low-fidelity polymerases, or tRNAs with reduced efficiency. As our analyses are limited to *in vitro* transcriptome data from the *Trichomonas* trophozoite stage under axenic culture conditions, it may be that the repertoire of RT genes in *Trichomonas* can be modulated under physiological or stress conditions or during the amoeboid or pseudocyst forms, similar to what is exhibited in different cell types and conditions of Dipteran flies.

## Conclusions

Our analysis of high-quality genome assemblies and long-read RNA-seq datasets from several *Trichomonas* species provides strong evidence that these parasites can selectively read through stop codons, thereby forming longer-than-predicted proteins. This is, to the best of our knowledge, the first time such a mechanism has been described in protozoan parasites. The RT genes characterized in *T. vaginalis* G3 encode 53 different protein functions, suggesting that a wide range of biological processes in the parasite can involve this mechanism. RT genes are physically located closer to each other than monocistronic genes, and generate among the most abundant transcripts by several orders of magnitude. The results presented in this manuscript are preliminary and will be updated with more accurate numbers and information on the mechanism the parasite uses to transcribe and translate these RT genes.

## Author contributions

F.C-H and J.M.C conceived and designed the study; M.S, F.C-H, and M.P undertook wet-lab work; S.A.S provided supervision; F.C-H performed bioinformatic analyses; F.C-H, J.M.C, and S.A.S wrote the original draft. All authors have read, revised, and approved the final version of the manuscript.

